# Characterization and functional analysis of *Toxoplasma* Golgi-associated proteins identified by proximity labelling

**DOI:** 10.1101/2024.02.02.578703

**Authors:** Rebecca R. Pasquarelli, Justin J. Quan, Emily S. Cheng, Vivian Yang, Timmie A. Britton, Jihui Sha, James A. Wohlschlegel, Peter J. Bradley

## Abstract

*Toxoplasma gondii* possesses a highly polarized secretory pathway that contains both broadly conserved eukaryotic organelles and unique apicomplexan organelles which play essential roles in the parasite’s lytic cycle. As in other eukaryotes, the *T. gondii* Golgi apparatus sorts and modifies proteins prior to their distribution to downstream organelles. Many of the typical trafficking factors found involved in these processes are missing from apicomplexan genomes, suggesting that these parasites have evolved unique proteins to fill these roles. Here we identify a novel Golgi-localizing protein (ULP1) which contains structural homology to the eukaryotic trafficking factor p115/Uso1. We demonstrate that depletion of ULP1 leads to a dramatic reduction in parasite fitness and replicative ability. Using ULP1 as bait for TurboID proximity labelling and immunoprecipitation, we identify eleven more novel Golgi-associated proteins and demonstrate that ULP1 interacts with the *T. gondii* COG complex. These proteins include both conserved trafficking factors and parasite-specific proteins. Using a conditional knockdown approach, we assess the effect of each of these eleven proteins on parasite fitness. Together, this work reveals a diverse set of novel *T. gondii* Golgi-associated proteins that play distinct roles in the secretory pathway. As several of these proteins are absent outside of the Apicomplexa, they represent potential targets for the development of novel therapeutics against these parasites.

**Importance:** Apicomplexan parasites such as *Toxoplasma gondii* infect a large percentage of the world’s population and cause substantial human disease. These widespread pathogens use specialized secretory organelles to infect their host cells, modulate host cell functions, and cause disease. While the functions of the secretory organelles are now better understood, the Golgi apparatus of the parasite remains largely unexplored, particularly regarding parasite-specific innovations that may help direct traffic intracellularly. In this work, we characterize ULP1, a protein that is unique to parasites but shares structural similarity to the eukaryotic trafficking factor p115/Uso1. We show that ULP1 plays an important role in parasite replication and demonstrate that it interacts with the conserved oligomeric Golgi (COG) complex. We then use ULP1 proximity labelling to identify eleven additional Golgi-associated proteins which we functionally analyze via conditional knockdown. This work expands our knowledge of the *Toxoplasma* Golgi apparatus and identifies potential targets for therapeutic intervention.

## Introduction

*Toxoplasma gondii* is an obligate intracellular parasite in the phylum Apicomplexa. This phylum includes parasites of both human and veterinary medical importance including *Plasmodium spp*. (malaria), *Cryptosporidium spp*. (diarrheal disease), *Eimeria spp*. (chicken coccidiosis), and *Neospora caninum* (neosporosis) (1–5). Approximately one-third of the global human population is chronically infected with *T. gondii* (6). While infection is usually asymptomatic in healthy individuals, the parasite can cause severe or fatal disease in immunocompromised people and congenitally infected neonates (7–9). While treatments exist that can suppress the acute infection, they are unable to clear the parasite completely resulting in lifelong chronic infection (8). A deeper understanding of apicomplexan biology is needed to inform the discovery of novel parasite-specific therapeutics.

Apicomplexan parasites possess a unique and highly polarized secretory pathway which plays an essential role in their life cycles (10,11). The endomembrane system of these parasites includes the endoplasmic reticulum and a Golgi apparatus, which is composed of a single stack of three to five cisternae, as well as several apicomplexan-specific organelles including the rhoptries, micronemes, dense granules, and inner membrane complex (IMC) (12,13). During parasite replication, these organelles are formed de novo from vesicles which bud from the Golgi and are trafficked through the secretory pathway to their destinations (14). The secretory pathway also delivers cargo to the plant-like vacuole (PLVAC), endosome like compartment (ELC), and a non-photosynthetic plastid called the apicoplast (15–18). As each of these organelles are essential, correct protein sorting is important for the survival of the parasite.

Recent studies have revealed a surprising intersection between *T. gondii*’s endocytic and exocytic pathways. Several Rabs which are components of the endosomal system in most eukaryotes, such as Rab5a and Rab7, have instead been repurposed as secretory trafficking proteins that reside in the ELC. The PLVAC, a hydrolytic compartment analogous to the lysosome, has been implicated in multiple roles such as autophagy, degradation of host-derived proteins, and proteolytic processing of proteins that are trafficked to the micronemes and rhoptries (16,19,20). Disruption of proteins that localize to the PLVAC and ELC or proteins that are typically involved in endosomal trafficking has been shown to cause defects in the biogenesis of rhoptries and micronemes (11,16,21–23). Furthermore, endolysosomal proteases have been shown to proteolytically process immature ROP and MIC proteins (16,19,20). Together, this evidence supports the convergence of endocytic and exocytic pathways in *T. gondii*.

Several key trafficking factors that play critical roles in protein trafficking in most eukaryotes are missing in apicomplexan genomes. Specifically, *T. gondii* and other apicomplexans lack most components of the Endosomal Sorting Complexes Required for Transport (ESCRT) and the Golgi-localized Gamma adaptin ear-containing ARF-binding (GGA) proteins (11,24,25). This limited repertoire of classical protein trafficking machinery suggests that apicomplexans may have evolved their own specialized proteins and pathways to facilitate these functions. Here we report the identification of a novel parasite-specific Golgi protein, ULP1, which is structurally homologous to the eukaryotic tethering factor p115/Uso1. We show that ULP1 plays a critical role in parasite fitness and its depletion results in defective parasite replication. We then use TurboID proximity labelling and immunoprecipitation to show that ULP1 interacts with the *T. gondii* conserved oligomeric Golgi (COG) complex and identify eleven previously uncharacterized Golgi-associated proteins. Finally, we analyze the orthology of each novel protein and use a conditional knockdown approach to assess their impact on parasite fitness, revealing several essential proteins and several more parasite-specific proteins.

## Results

### TGGT1_289120 is a parasite-specific Golgi protein that is important for parasite fitness

We identified a 290 kDa hypothetical protein with the gene ID TGGT1_289120 during our previous proximity labelling experiments using IMC proteins as bait (26). BLASTp analysis revealed that TGGT1_289120 has orthologs in *Hammondia*, *Neospora*, *Besnoitia, Cystoisospora*, and *Sarcocystis* but appears to be absent from other apicomplexan parasites and higher eukaryotes (27). Secondary structure analysis using Phyre2 revealed that residues 495-647 of TGGT1_289120 contain structural homology to the globular head domain of p115/Uso1, a membrane tethering protein conserved in eukaryotes (28,29). InterPro also identified an armadillo (ARM)-like domain within this region, and DeepCoil2 predicted a coiled-coil (CC) domain within residues 1736-1812 (Fig 1A) (30,31). p115/Uso1 also contains ARM repeats within its globular head domain and a single CC domain near the C-terminus, further supporting their structural similarity (29). While this study was being prepared for publication, a study of the *T. gondii* COG complex by Marsilia et al. independently identified TGGT1_289120 and named it Uso1-like protein 1 (TgULP1) based on its structural similarity to the p115/Uso1 protein (32). To maintain consistency, we adopted the same nomenclature and also named the protein ULP1.

**Fig 1.**
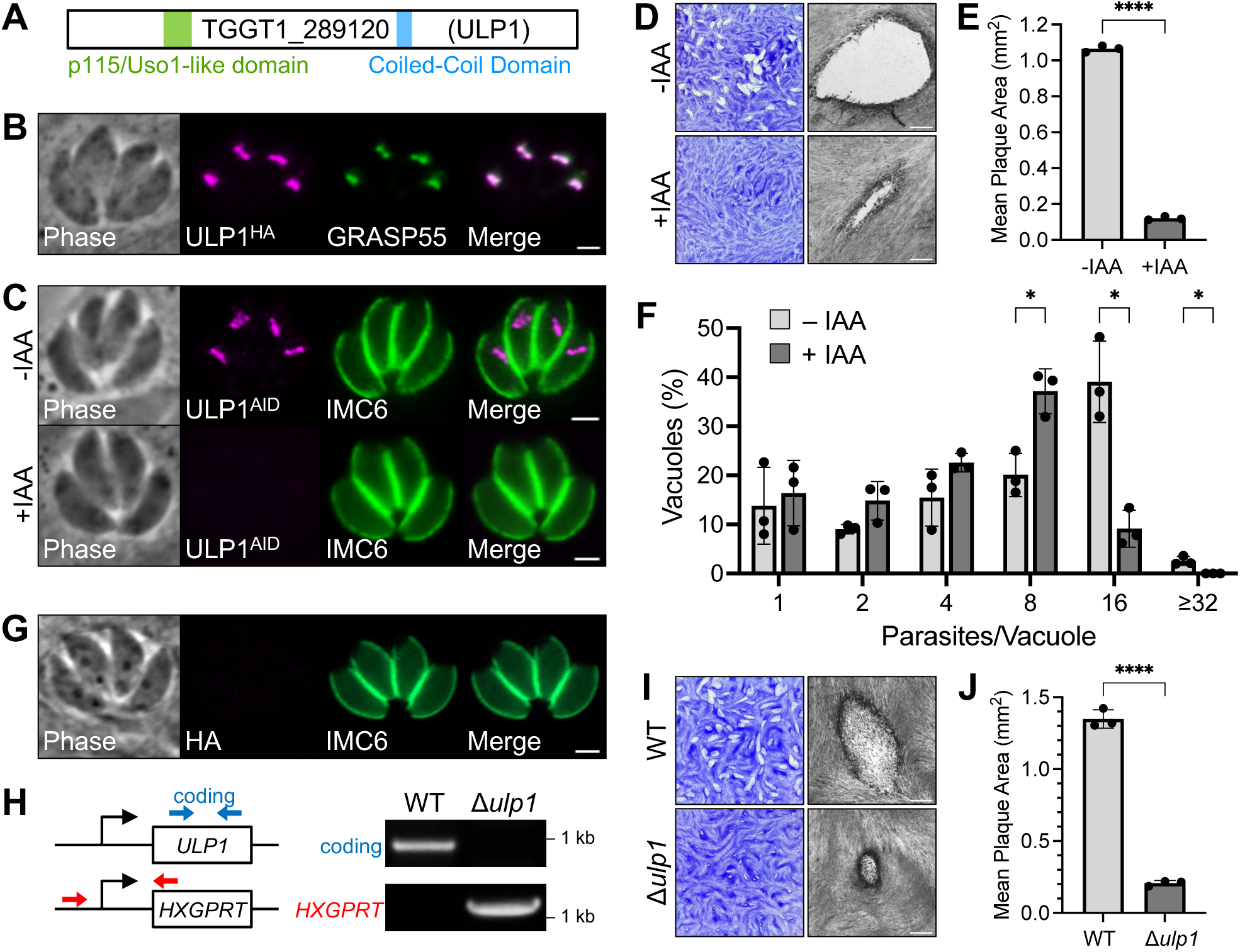
ULP1 is a novel Golgi-localizing protein that is important for parasite fitness. A) Gene model of TGGT1_289120 (ULP1) showing its p115/Uso1-like domain and predicted CC domain. B) IFA showing that ULP1 colocalizes with the Golgi apparatus marker GRASP55-YFP. Magenta = anti-HA detecting ULP1^3xHA^, Green = GRASP55-YFP. C) IFA showing that ULP1^AID^ localizes normally and is depleted after 24 hours of IAA treatment. Magenta = anti-HA detecting ULP1^AID^, Green = anti-IMC6. D) Plaque assay for ULP1^AID^ parasites -/+IAA shows that depletion of ULP1 results in a severe reduction in overall lytic ability. E) Quantification of plaque size for plaque assays shown in panel D. Significance was determined using two-tailed t test. **** = P < 0.0001. F) Quantification of the number of parasites per vacuole for ULP1^AID^ parasites treated with IAA or vehicle control after 24 hours of growth. Significance was determined using multiple two-tailed t tests. * = P < 0.05. G) IFA of Δ*ulp1* parasites confirms loss of ULP1^3xHA^ signal. Magenta = anti-HA, Green = anti-IMC6. H) PCR verification for genomic DNA of WT and Δ*ulp1* parasites. Diagram indicates the binding location of primers used to amplify the ULP1 coding sequencing (blue arrows) and the site of recombination for the knockout (red arrows). I) Plaque assays of WT and Δ*ulp1* parasites. J) Quantification of plaque size for plaque assays shown in panel I. Statistical significance was determined using a two-tailed t test. **** = P < 0.0001. Scale bars for plaque assays = 0.5 mm. Scale bars for IFAs = 2 µm.

The similarity to the essential vesicular trafficking factor p115/Uso1 and its low phenotype score of -3.8 in a genome wide CRISPR/Cas9 screen (GWCS) led us to hypothesize that this protein may play a key role in secretory trafficking in *T. gondii* (33). To determine the localization of ULP1, we used CRISPR/Cas9 to fuse sequences encoding a C-terminal 3xHA epitope tag at its endogenous locus (34,35). Immunofluorescence assay (IFA) revealed that ULP1 localized to a distinct bar-shaped region anterior to the nucleus, suggesting localization to the Golgi apparatus. To confirm this, we co-stained for GRASP55-YFP, a marker of the cis-medial Golgi apparatus in *T. gondii* and found that the two proteins colocalized (Fig 1B) (36).

To study the function of ULP1, we used the auxin-inducible degron (AID) system which allows rapid proteasomal degradation of a target protein upon treatment with indoleacetic acid (IAA) (37,38). To create an ULP1^AID^ parasite strain, we fused sequences encoding an mAID-3xHA tag to the 3’ end of the gene in a strain that carries the TIR1 auxin-receptor FBOX protein. IFA showed that the degron-tagged protein localizes appropriately and is completely depleted after 24 hours of IAA treatment (Fig 1C). The ULP1-depleted parasites did not appear to exhibit defects in overall morphology. To assess how loss of ULP1 would affect the parasite’s overall lytic ability, we performed plaque assays, which showed a dramatic 88.7% decrease in plaque size when ULP1 was depleted (Fig 1D, E). To determine whether this defect in lytic ability was due to defects in replication, we quantified the number of parasites per vacuole after 30 hours of growth for parasites treated with IAA or a vehicle control. At 30 hours post-infection, ULP1-depleted parasites were found primarily in 8-parasite vacuoles, whereas untreated parasites were primarily found in 16-parasite vacuoles (Fig 2F). This indicates that the defect in overall lytic ability is due to a defect in replication. Given the low phenotype score (-3.8), we wondered whether the slow growth and ability to form small plaques may be due to some residual ULP1 which escaped degradation. To determine if this was the case, we used CRISPR/Cas9 to disrupt the endogenous locus for ULP1 and verified loss of the protein by both IFA and PCR (Fig 1G-H). Using this method, we successfully generated a Δ*ulp1* strain which phenocopied the plaque defect exhibited by IAA-treated ULP1^AID^ parasites (Fig 1I-J). To avoid potential issues with functional compensation, which has been previously documented for knockout strains in *T. gondii*, we decided to proceed with the ULP1^AID^ strain for further phenotypic analysis (39–41).

**Fig 2.**
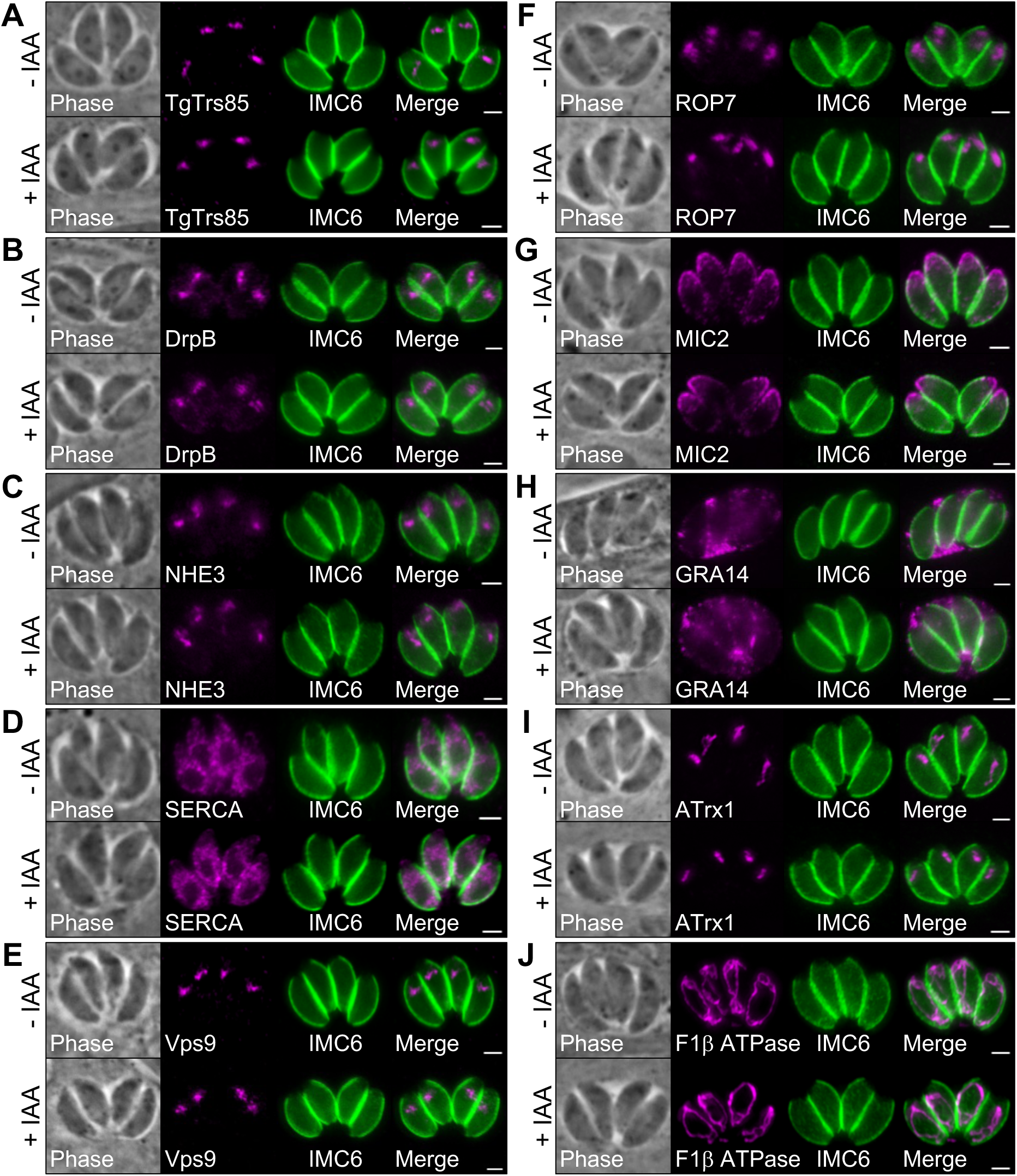
Depletion of ULP1 does not affect overall organellar morphology. IFAs comparing the overall morphology of various *T. gondii* organelles in control vs. ULP1-depleted parasites. A) Golgi apparatus morphology is unaffected by depletion of ULP1. Magenta = anti-V5 detecting TgTrs85^3xV5^, Green = anti-IMC6. B) Post-Golgi vesicle morphology is unaffected by depletion of ULP1. Magenta = anti-DrpB, Green = anti-IMC6. C) PLVAC morphology is unaffected by depletion of ULP1. Magenta = anti-NHE3, Green = anti-IMC6. D) ER morphology is unaffected by depletion of ULP1. Magenta = anti-SERCA, Green = anti-IMC6. E) ELC morphology is unaffected by depletion of ULP1. Magenta = anti-V5 detecting Vps9^3xV5^, Green = anti-IMC6. F) Rhoptry morphology is unaffected by depletion of ULP1. Magenta = anti-ROP7, Green = anti-IMC6. G) Microneme morphology is unaffected by depletion of ULP1. Magenta = anti-MIC2, Green = anti-IMC6. H) Dense granule morphology is unaffected by depletion of ULP1. Magenta = anti-GRA12, Green = anti-IMC6. I) Apicoplast morphology is unaffected by depletion of ULP1. Magenta = anti-ATrx1, Green = anti-IMC6. J) Mitochondrion morphology is unaffected by depletion of ULP1. Magenta = anti-F1β ATPase, Green = anti-IMC6. Scale bars = 2 µm.

Next, we used IFA to assess whether depletion of ULP1 resulted in aberrant organellar morphology. We began by endogenously tagging the known Golgi protein TgTrs85 to determine whether loss of ULP1 would affect the Golgi apparatus (18). When we depleted ULP1, we found that TgTrs85 was unaffected (Fig 2A). We next stained for DrpB, a dynamin-related protein that localizes to post-Golgi vesicles, which also appeared to be unaffected (Fig 2B) (23). Other components of the secretory pathway such as the PLVAC (marked by NHE3), endoplasmic reticulum (SERCA), and ELC (Vps9) were similarly unaffected (Fig 2C-E). Next, we wanted to determine whether the morphology of downstream secretory organelles was altered, so we stained for the rhoptries (ROP7), micronemes (MIC2), and dense granules (GRA14). We additionally stained for the apicoplast (Atrx1) and mitochondrion (F1β ATPase). Surprisingly, we did not observe mislocalization of any of these markers, indicating these organelles are not grossly affected upon ULP1 knockdown (Fig 2F-J).

### ULP1 proximity labelling and immunoprecipitation yields candidate Golgi proteins

To identify additional novel Golgi proteins, we decided to use ULP1 as bait for both proximity labeling and immunoprecipitation (IP) experiments. For proximity labelling, the biotin ligase TurboID-3xHA was fused to the C-terminus of ULP1 (Fig 3A) (42). The ULP1^TurboID^ fusion protein was detectable by IFA and trafficked appropriately to the Golgi apparatus. In the absence of biotin, only the endogenously biotinylated apicoplast stained with streptavidin. Upon addition of biotin, the Golgi apparatus also stained with streptavidin, confirming that the ULP1^TurboID^ fusion protein was enzymatically active and biotinylated proximal Golgi proteins (Fig 3B). Western blot analysis confirmed that the addition of biotin resulted in dramatically increased amounts of biotinylated proteins in ULP1^TurboID^ lysates (Fig 3C). The IP analysis was performed on the ULP1^3xHA^ strain. Western blot showed that the 3xHA-tagged protein migrated at a smaller size than expected based on its molecular weight of 290 kDa and exhibited significant protein breakdown. During the process of affinity chromatography for the IP experiment, the protein experienced even more severe breakdown. Despite this, the elution fraction was still highly enriched for the HA-tagged protein (Fig 3D). Thus, we continued with the experiment.

**Fig 3.**
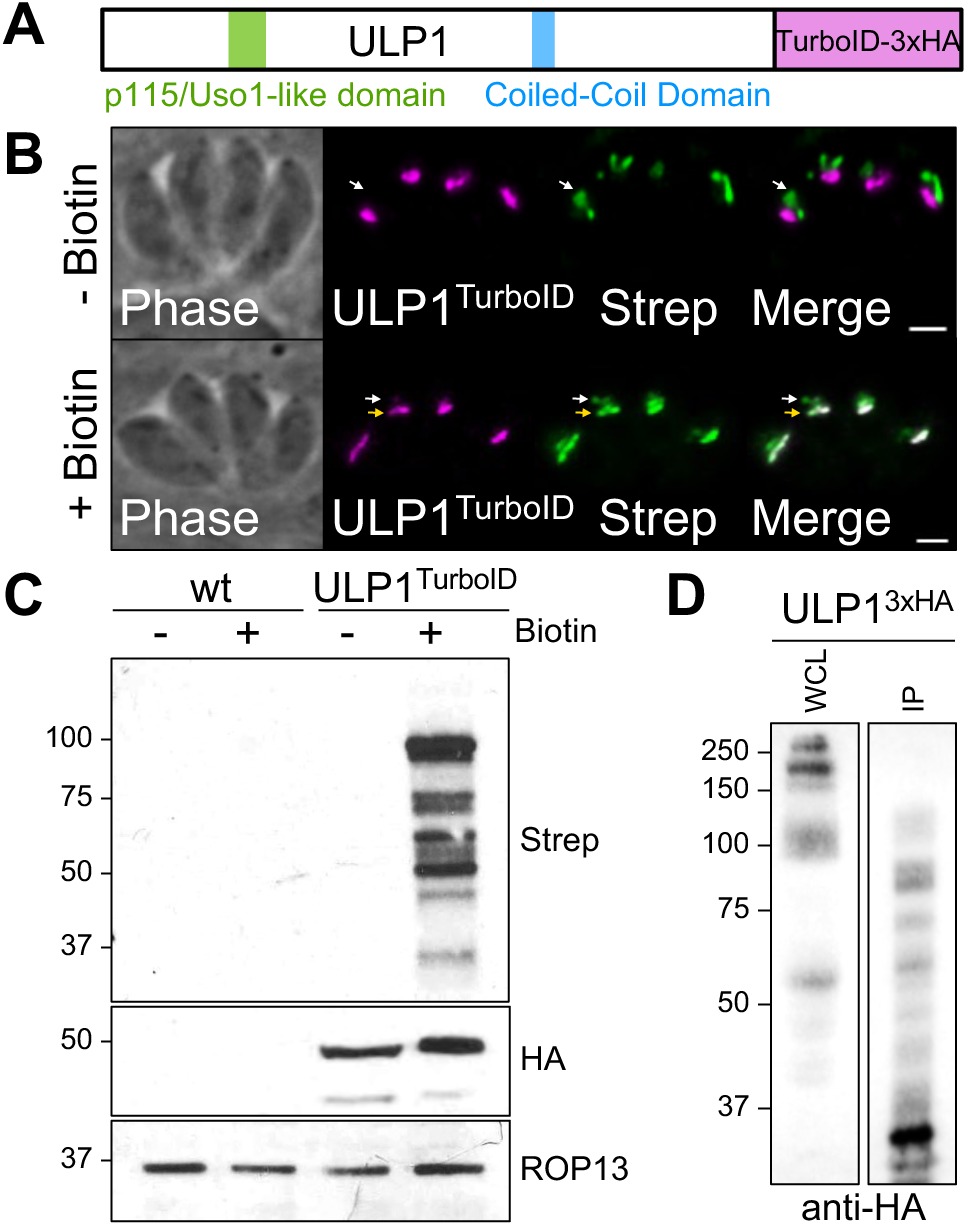
Validation of ULP1 TurboID and IP. A) The TurboID biotin ligase and a 3xHA tag were fused to the C-terminus of ULP1 to generate the ULP1^TurboID^ line. B) IFA showing that ULP1^TurboID^ localizes as expected and results in biotinylation of proximal proteins in a biotin-dependent manner (yellow arrow). The apicoplast, which contains endogenously biotinylated proteins, is denoted by white arrow. Magenta = anti-HA detecting ULP1^TurboID^, Green = streptavidin. Scale bar = 2 µm. C) Western blot showing biotin-dependent biotinylation of proteins in the ULP1^TurboID^ line, which is absent from a wild-type control. ROP13 is used as a loading control. D) Western blot showing enrichment of ULP1^3xHA^ after immunoprecipitation with anti-HA resin despite significant protein breakdown. WCL = whole-cell lysate prior to IP. IP = final sample eluted from anti-HA resin.

As expected, the bait protein ULP1 was the top hit identified by mass spectrometry in the IP experiments and was among the top 5 hits in the TurboID experiment (Table S1 and Table S2). DrpB, TgTrs85, TgEpsL, and Vps9, which have all been previously shown to localize to the *Toxoplasma* Golgi apparatus or nearby compartments of the secretory pathway, were highly enriched in our TurboID experiment (Table S1) (18,23,43,44). In addition, many of the top hits are predicted to be involved in processes such as ER-to-Golgi retrograde transport, Golgi organization, and intracellular protein transport. Many of the most enriched proteins were hypothetical proteins that have not previously been studied. Domain analysis allowed us to identify two of these hypothetical proteins, TGGT1_278890 and TGGT1_262450, as homologs of the known Golgi proteins CDP/cut alternatively spliced product (CASP) and dymeclin (DYM), respectively.

Our IP experiment identified several interesting candidate interactors, including the hypothetical protein TGGT1_216370, which was also among the top hits in the TurboID experiment (Table S2). In addition, homologs of all eight subunits of the COG complex immunoprecipitated with ULP1 and were highly enriched in our TurboID experiment. All eight are currently annotated as hypothetical proteins, but BLASTp and functional domain analysis supports their designation as COG complex subunits, although the BLASTp e-value for COG1 is weak (e^-6^) (24). As p115/Uso1 is a known interactor of the COG complex via direct binding to COG2, this result further supports the designation of ULP1 as a p115/Uso1-like protein (45). Several additional proteins were found to immunoprecipitate with ULP1, although fewer peptides were recovered for these proteins. Among these lower abundance hits were a Sec23/Sec24 trunk domain-containing protein, TgEpsL, COP1 subunit ε, the cyclin-dependent kinase (CDK)-related kinase Crk2, Vps35, Rab7, AP1-γ, and DrpB.

### Verification of candidate Golgi proteins

To continue our study of the *Toxoplasma* Golgi apparatus, we selected eleven uncharacterized proteins with GWCS phenotype scores of less than -2 from our TurboID and IP hits (Table 1). Candidates were prioritized based on functional domain analysis and orthology to known Golgi proteins. Five of the selected genes appeared to be direct orthologs of known Golgi-associated proteins. TGGT1_290310 is the putative COG1 orthologue that we identified in our ULP1 IP. TGGT1_311400 is annotated as WD domain, G-beta repeat-containing protein, but BLASTp analysis revealed its identity as Sec31. TGGT1_232190 contains a Sec7 domain within a larger Golgi Brefeldin A resistance guanine nucleotide exchange Factor 1 (GBF1) domain. TGGT1_264090 contains a TRAPPC11 domain. TGGT1_301410 contains both an ENTH domain and a Tepsin domain. Two proteins contain functional domains that suggested they may have functions related to the secretory system, but do not appear to be direct orthologs of any known Golgi proteins. TGGT1_294730 contains an ancestral coatomer element 1 (ACE1) domain from Sec16/Sec31 yet lacked other key domains found in these proteins. TGGT1_230400 contains a WIP-related protein domain but appears to only have orthologs in other coccidian parasites (46). The final four selected genes (TGGT1_216370, TGGT1_207370, TGGT1_258080, and TGGT1_240220) are hypothetical proteins that contained no identifiable functional domains aside from predicted CC domains and had orthologs only in cyst-forming coccidia.

**Table 1.** Summary of Golgi-associated proteins identified in this study. Table summarizing the twelve novel Golgi-associated proteins identified and characterized in this study, listed in order of enrichment in the ULP1 TurboID experiment. Average spectral counts for each protein in the TurboID and IP experiments are shown (“Ctrl” = control, “Exp” = experimental). GWCS = phenotype score assigned in a genome wide CRISPR/Cas9 screen (33). Abbrv. = protein name abbreviation.

| Gene ID | Abbrev. | Protein Name | TurboID |  | IP |  | ToxoDB Annotation | GWCS |
| --- | --- | --- | --- | --- | --- | --- | --- | --- |
|  |  |  | Ctrl | Exp | Ctrl | Exp |  |  |
| TGGT1_289120 | ULP1 | Uso1-Like Protein 1 | 1.5 | 341 | 0 | 382 | hypothetical protein | -3.8 |
| TGGT1_311400 | Sec31 | Secretory 31 | 28 | 698.5 | 0 | 0 | WD domain, G-beta repeat-containing protein | -5.53 |
| TGGT1_294730 | Sec16-L | Sec16-like Protein | 12 | 468 | 0 | 0 | hypothetical protein | -2.21 |
| TGGT1_264090 | TRAPPC11 | Trafficking protein particle complex subunit 11 | 2 | 326.5 | 0 | 0 | hypothetical protein | -4.74 |
| TGGT1_290310 | COG1 | Conserved Oligomeric Golgi complex component 1 | 0 | 199 | 0 | 59 | hypothetical protein | -3.75 |
| TGGT1_232190 | GBF1 | Golgi Brefeldin A resistance guanine nucleotide exchange Factor 1 | 1 | 157 | 0 | 0 | Sec7 domain-containing protein | -5.57 |
| TGGT1_207370 | GLP2 | Golgi Localizing Protein 2 | 1 | 126.5 | 0 | 0 | hypothetical protein | -3.89 |
| TGGT1_230400 | WPDP | WIP-related Protein Domain-containing Protein | 2 | 67 | 0 | 0 | hypothetical protein | -2.42 |
| TGGT1_216370 | GLP3 | Golgi Localizing Protein 3 | 0 | 46 | 0 | 38 | hypothetical protein | -4.37 |
| TGGT1_301410 | Tepsin | AP-4 complex accessory subunit Tepsin | 0 | 41 | 0 | 0 | hypothetical protein | -4.16 |
| TGGT1_258080 | GLP1 | Golgi Localizing Protein 1 | 0 | 32.5 | 0 | 0 | hypothetical protein | -2.41 |
| TGGT1_240220 | GLP4 | Golgi Localizing Protein 4 | 0 | 20 | 0 | 0 | hypothetical protein | -2.1 |

To verify these candidate Golgi proteins, we endogenously tagged each protein at its C-terminus with an auxin-inducible degron coupled to a 3xHA epitope tag in a TIR1 strain (37). We then expressed the Golgi marker GRASP55-YFP in each line and used IFA to assess the localization of each gene (36). All eleven genes either colocalized with GRASP55 or were closely adjacent, confirming their association with the *Toxoplasma* Golgi apparatus (Fig 4). Based on their localization and functional domains, we assigned names to each of the eleven proteins.

**Fig 4.**
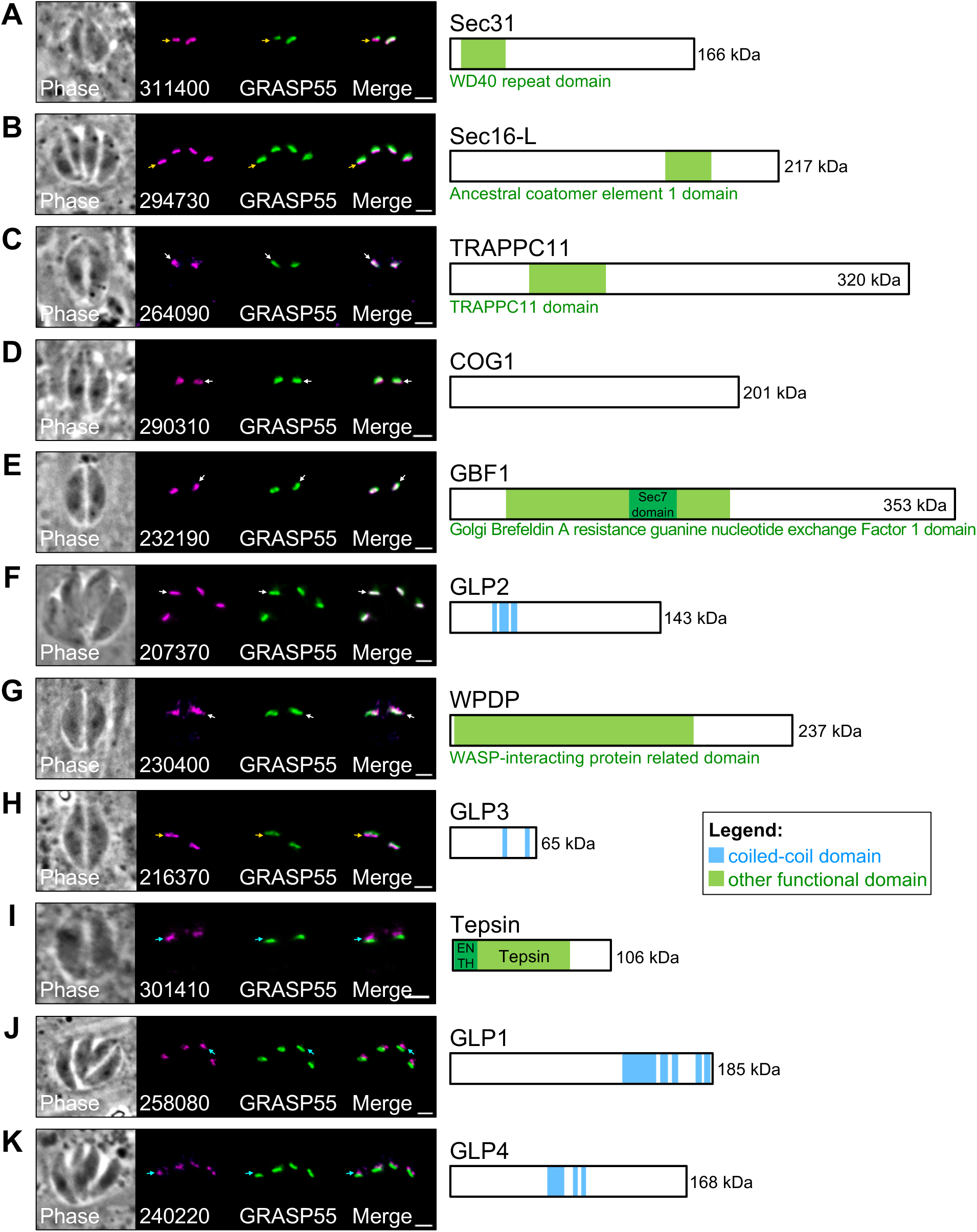
ULP1 TurboID and IP hits reveals novel Golgi-associated proteins. IFAs and corresponding gene models for 11 genes identified as candidate Golgi proteins in the ULP1 TurboID and IP experiments. Each gene was endogenously tagged with mAID-3xHA or mIAA7-3xHA auxin-inducible degrons. White arrows indicate that the protein primarily colocalizes with GRASP55-YFP. Yellow arrows indicate that the protein primarily localizes upstream of GRASP55-YFP. Cyan arrows indicate that the protein primarily localizes downstream of GRASP55-YFP. A) TGGT1_311400 (Sec31) localizes upstream of GRASP55 and contains a WD40 repeat domain. B) TGGT1_294730 (Sec16-L) localizes upstream of GRASP55 and contains an ACE1 domain. C) TGGT1_264090 (TRAPPC11) colocalizes with GRASP55 and contains a TRAPPC11 domain. D) TGGT1_290310 (COG1) colocalizes with GRASP55 and contains no identifiable functional domains. E) TGGT1_232190 (GBF1) colocalizes with GRASP55 and contains a GBF1 domain. F) TGGT1_207370 (GLP2) colocalizes with GRASP55 and contains three predicted CC domains within residues 273-301, 313-373, and 382-423. G) TGGT1_230400 (WPDP) colocalizes with GRASP55 and contains a large WASP-interacting protein related domain. H) TGGT1_216370 (GLP3) localizes downstream of GRASP55 and contains two predicted CC domains within residues 337-370 and 494-527. I) TGGT1_ 301410 (Tepsin) localizes downstream of GRASP55 and contains ENTH and Tepsin domains. J) TGGT1_258080 (GLP1) localizes downstream of GRASP55 and contains five predicted CC domains within residues 1117-1332, 1362-1407, 1425-1464, 1603-1646, and 1659-1701. K) TGGT1_240220 (GLP4) localizes downstream of GRASP55 and contains three predicted CC domains within residues 634-747, 797-830, and 850-881. Magenta = anti-HA detecting degron-tagged proteins, Green = GRASP55-YFP. Scale bars = 2 µm.

TGGT1_311400 was designated Sec31, TGGT1_294730 as Sec16-like protein (Sec16-L), TGGT1_264090 as TRAPPC11-like protein (TRAPPC11), TGGT1_290310 as COG1, TGGT1_232190 as GBF1, and TGGT1_230400 as WIP-related Protein Domain-containing Protein (WPDP). The hypothetical protein TGGT1_258080 was also recently identified by Marsilia et al., so we adopted their nomenclature and named it Golgi Localizing Protein 1 (GLP1). The hypothetical proteins TGGT1_216370, TGGT1_207370, TGGT1_240220 were designated as Golgi Localizing Protein 2, 3, and 4, respectively (GLP2-4). Upon further inspection, we noticed that the extent to which each protein colocalized with GRASP55 varied. Sec31, Sec16-L, and GLP3 appeared to localize upstream of GRASP55 in the secretory pathway (Fig 4A, B, and H). Sec31 and Sec16 are both known to localize to ER exit sites where they play roles in anterograde transport (47). While GLP3 does not contain any domains that hint at its function, its similar localization to Sec31 and Sec16-L suggests it may also play a role in early anterograde transport. TRAPPC11, COG1, GBF1, GLP2, and WPDP all appeared to mostly colocalize with GRASP55, indicating a primary localization in the cis-medial Golgi (Fig 4C-G and I). TRAPPC11 and WPDP displayed some additional signal downstream of GRASP55, suggesting they may also localize to the trans Golgi network (Fig 4C and G). Finally, Tepsin, GLP1, and GLP4 localized downstream of GRASP55, indicating that they likely are restricted to the trans Golgi network (Fig 4J and K).

### Functional analysis of novel proteins

We next used the AID system to assess how depletion of each novel Golgi protein would affect parasite fitness. Five of the eleven strains (Sec31, GBF1, COG1, Tepsin, and TRAPPC11) were found to be completely deficient in plaque formation, indicating that these proteins are essential for parasite survival (Fig 5A). Sec31 exhibited a severe growth arrest, with nearly all vacuoles containing only a single parasite after 24 hours of growth in the presence of IAA (Fig 5B). Despite the severe growth defect, Sec31-depleted parasites appeared to have normal overall morphology as assessed by IMC6 staining. GBF1 also exhibited a severe growth arrest (Fig 5C). However, in contrast to Sec31, many GBF1-depleted parasites also exhibited defective IMC morphology. Both COG1 and Tepsin-depleted parasites did not exhibit growth arrest at 24 hours but had severe defects in morphology, such as large gashes in the cytoskeleton marked by IMC6 (Fig 5D, E). The last essential protein, TRAPPC11, did not exhibit growth arrest or any apparent defects in overall morphology as assessed by IMC6 staining after 24 hours of IAA treatment (Fig 5F).

**Fig 5.**
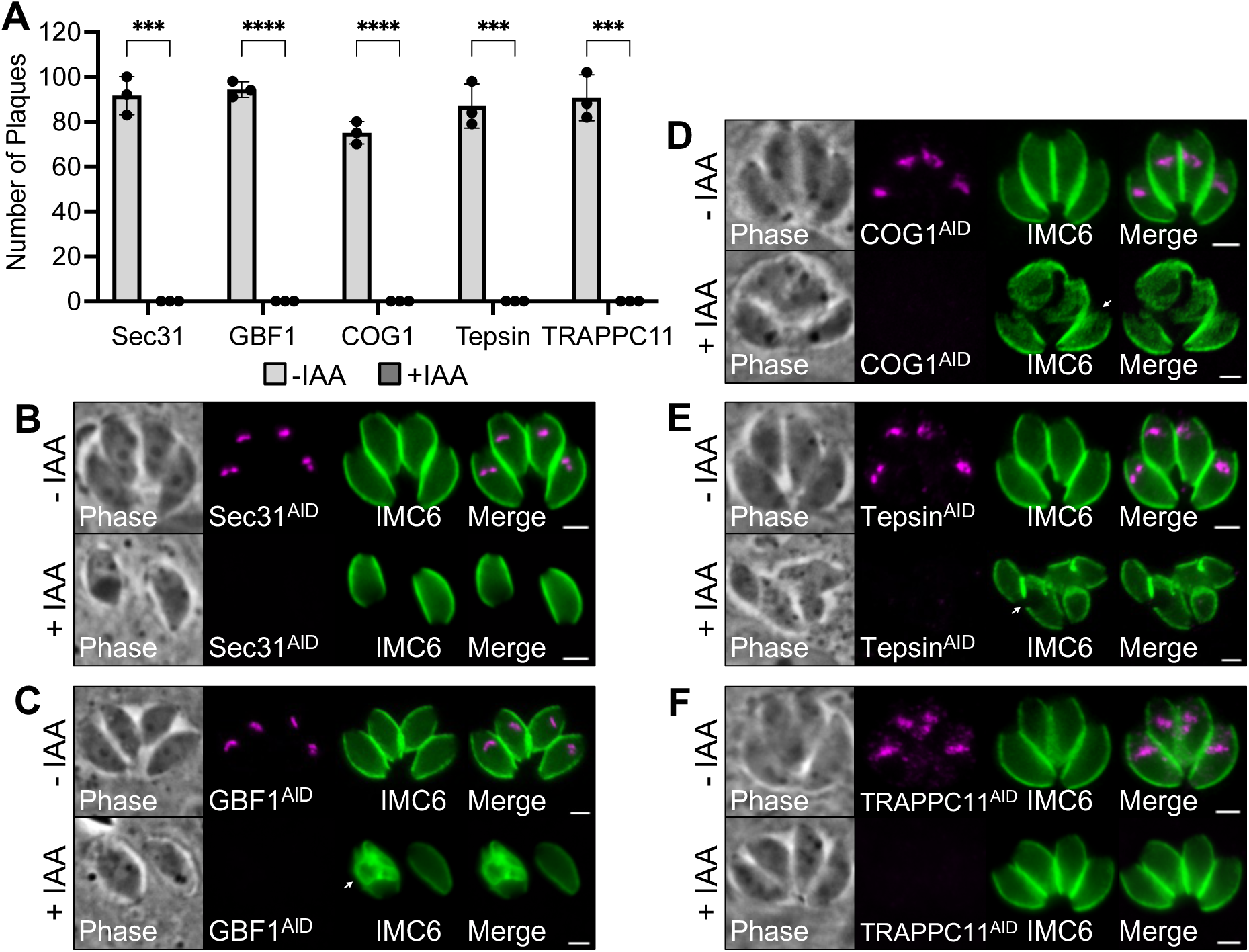
Conditional knockdown of Sec31, GBF1, COG1, Tepsin, and TRAPPC11. A) Plaque assays were performed -/+ IAA to assess how depletion of Sec31, GBF1, COG1, Tepsin, or TRAPPC11 affects overall lytic ability over the course of seven days. Number of plaques was quantified for each condition. Statistical significance was determined using multiple two-tailed t tests (**** = P < 0.0001, *** = P < 0.001) B, C) Depletion of Sec31 or GBF1 results in growth arrest. Depletion of GBF1 additionally leads to morphological defects in some parasites (white arrow). C, D) Depletion of COG1 or Tepsin results in morphological defects (white arrows). F) Depletion of TRAPPC11 does not result in any obvious defects in morphology. Magenta = anti-HA detecting degron-tagged proteins, Green = anti-IMC6. Scale bars = 2 µm.

The remaining six proteins were found by plaque assay to be non-essential, but their depletion resulted in a statistically significant reduction in lytic ability (Fig 6A). The severity of the defect varied widely among the six proteins. The most severe growth defect (91.8%) was observed for GLP2. The remaining five proteins exhibited more modest reductions – 25.7% for Sec16-L, 21.0% for WPDP, 14.2% for GLP1, 40.5% for GLP3, 37.9% for GLP4. Despite the reduction in fitness, all six of these proteins exhibited no obvious defects in morphology by IFA (Fig 6B-G).

**Fig 6.**
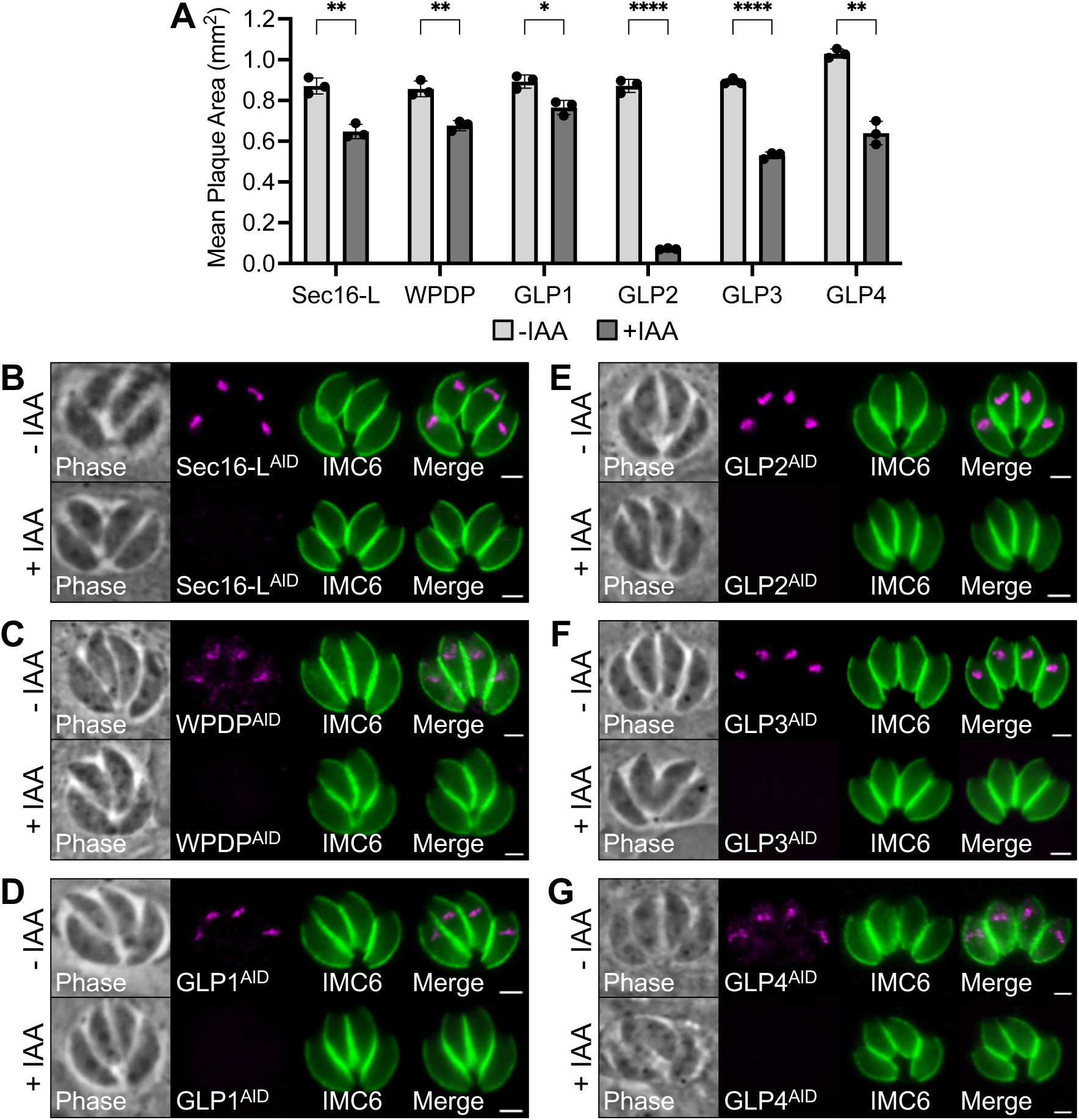
Conditional knockdown of Sec16-L, WPDP, GLP1, GLP2, GLP3, and GLP4. A) Plaque assays were performed -/+IAA to assess how depletion of Sec16-L, WPDP, GLP1, GLP2, GLP3, and GLP4 affects overall lytic ability over the course of seven days. Mean plaque area was quantified for each condition. Statistical significance was determined using multiple two-tailed t tests (**** = P < 0.0001, *** = P < 0.001, ** = P < 0.01, * = P <0.05.) B-G) Depletion of Sec16-L, WPDP, GLP1, GLP2, GLP3, or GLP4 does not result in any obvious defects in morphology. Magenta = anti-HA detecting degron-tagged proteins, Green = anti-IMC6. Scale bars = 2 µm.

## Discussion

In this study, we identified and characterized the novel Golgi protein ULP1 as a critical regulator of parasite growth and used it as a probe to identify and characterize additional novel Golgi proteins. Our analysis of ULP1 demonstrated that the protein localizes to the cis-medial Golgi and plays an important role in parasite replication. Loss of the protein by either AID-mediated knockdown or gene knockout resulted in a severe defect in plaque size. Our findings contrast with that of Marsilia et al. who showed a complete loss of lytic ability upon ULP1 knockdown and place its primary localization in the trans-Golgi network (32). While the conditional knockdown strains may vary in the phenotypes observed, the ULP1 knockout indicates that the protein is not strictly required for parasite survival. ULP1 is an interesting protein because it is unique to cyst-forming coccidia yet is structurally homologous to the conserved eukaryotic trafficking factor p115/Uso1. Studies in mammalian cells, *Drosophila*, and yeast have demonstrated that p115/Uso1 has a wide range of essential functions including tethering COPI vesicles to the Golgi, transporting vesicles from the ER to the Golgi, and facilitating SNARE complex assembly (48–53). Loss of p115/Uso1 has been shown to result in fragmentation of the Golgi and accumulation of Golgi-derived vesicles (54). In addition to its localization to the Golgi apparatus, the protein has been observed associating with the spindle poles, and loss of the protein has been shown to impair mitosis (55,56). While we did not identify any apparent defects in trafficking of markers to key secretory organelles or loss of Golgi integrity upon ULP1 knockdown, we did observe a significant reduction in replicative ability which could suggest that ULP1 plays a role in cell cycle regulation, similar to p115/Uso1. P115/Uso1 is known to interact with several trafficking proteins including Rab1, GBF1, COG2, GM130, and SNARE complex components Syntaxin 5, Bet1 and Bos1 (45,51,57–59). Our IP analysis was highly enriched for all eight COG complex subunits, with COG2 being the second most enriched. Marsilia et. al also identified all eight components of the COG complex in their ULP1 IP, further supporting the legitimacy of these interactions (32). Interestingly, we did not identify any of the other known p115/Uso1 interacting proteins. While these other proteins may have been lost during processing, it is also possible that ULP1 function has diverged enough from p115/Uso1 such that it does not bind to these proteins and instead uses other parasite-specific proteins to fulfill these roles. Supporting this, the novel Golgi protein GLP2 was found to IP with ULP1 in both Marsilia et. al and our study. p115/Uso1 has been shown to bind to Cdk1 and loss of p115 was demonstrated to reduce Cdk1 activation (56). Interestingly, we identified a CDK-related kinase, Crk2, in our IP analysis. Given that ULP1 depletion leads to a decrease in replication rate, it is possible that interaction with Crk2 plays a role in *T. gondii* mitosis. Alternatively, the replication defect could be due to general ER stress caused by alteration of the secretory pathway (60). When comparing the remaining results of our IP with Marsilia et. al, several differences are observed. They identified a wide variety of other proteins including eight SNARE proteins, four AP-5 complex proteins, and five COPI complex subunits that were either absent from our analysis or did not reach the two-fold enrichment cut-off. Additionally, the authors noted that over 20 IMC proteins and Rab11b, which is critical for IMC biogenesis, were enriched in their ULP1 IP. However, we did not observe enrichment for any of these proteins. Similarly, our IP identified Crk2, Rab7, Vps35, TgEpsL, AP1-γ, and a Sec23/Sec24 trunk domain-containing protein while the other study did not. These disparities likely arise from differences in sample processing, and further analysis of ULP1 binding partners will be needed to create a more definitive picture of the ULP1 interactome.

Using ULP1 as bait in both TurboID and IP experiments, we were able to identify 11 previously uncharacterized Golgi-associated proteins. Sec31, COG1, GBF1, Tepsin, and TRAPPC11 were found to be essential, which is unsurprising due to their critical conserved functions in other eukaryotes. Two of these proteins, Sec31 and GBF1, caused an immediate growth arrest upon depletion, likely due to a block in the secretory pathway that prevents the continuation of the cell cycle. Sec31 acts as a component of the COPII complex, which facilitates anterograde transport from the ER to the Golgi and causes growth arrest when depleted in *Trypanosomes* (47,61,62). We observed the *T. gondii* Sec31 localizing upstream of GRASP55 in the secretory pathway, indicating that it likely localizes to ER exit sites as in other eukaryotes. The localization of this protein to this subcompartment of the Golgi represents a valuable new marker for this region of the secretory pathway in *T. gondii*. GBF1 is a Golgi-resident guanine nucleotide exchange factor that regulates COPI vesicle-mediated transport by activating ADP-ribosylation factor (Arf) proteins (63,64). We observed *Toxoplasma* GBF1 localizing to the cis-medial Golgi, which agrees with its function in other systems.

Unlike Sec31 and GBF1, the remaining three essential proteins, COG1, Tepsin, and TRAPPC11, did not exhibit an immediate growth arrest upon depletion. COG1 is a component of the COG complex, which acts in the retrograde transport pathway by tethering COPI-coated vesicles that are used for recycling Golgi-resident enzymes such as glycosyltransferases back to the Golgi (65,66). Several *T. gondii* proteins such as GAP50 and CST1 have been shown to be glycosylated, and this modification is required for their proper targeting and function (67,68). In agreement with this, Marsilia et. al demonstrated that O-glycosylation is disrupted upon COG3 or COG7 depletion (32). This function is primarily carried out in the cis/medial Golgi, which matches our observed colocalization of COG1 with GRASP55. Tepsin is an accessory protein that associates with the AP-4 complex, which plays a role in vesicular trafficking of proteins at the trans-Golgi network (69,70). Knockdown of either COG1 or Tepsin resulted in severe defects in IMC morphology, suggesting that loss of either protein causes issues with the secretory pathway ultimately leading to defects in IMC biogenesis. TRAPPC11, on the other hand, did not exhibit any obvious morphological defects. TRAPPC11 was observed colocalizing with GRASP55 in the cis-medial Golgi. Since TRAPPC11 functions in anterograde vesicle transport as a member of the TRAPPIII tethering complex, it is possible that *Toxoplasma* TRAPPC11 could play a key role in trafficking of effector proteins like ROPs and MICs, leading to a defect in invasion and/or egress (71). Further studies will be needed to assess precisely how each of these essential proteins affects parasite fitness.

Two of the identified proteins, Sec16-L and WPDP, do not appear to be direct orthologs of known Golgi proteins but contain domains that suggested conserved functions. Sec16-L contains the ACE1 domain from Sec16, an essential COPII vesicle coat protein required for ER transport vesicle budding and formation of ER exit sites (72). However, it is missing the Sec16 conserved C-terminal domain which facilitates binding to Sec23 and is essential for function (73). Sec16-L’s localization upstream of GRASP55 further supports its functional similarity to Sec16 and provides another marker of this early subcompartment of the secretory pathway in addition to Sec31. While Sec16 is essential in other systems, Sec16-L had a relatively modest fitness defect upon conditional knockdown and its ortholog in *P. falciparum* (PF3D7_1119900) is also predicted to be dispensable (74). It is possible that another protein has adopted this function in *T. gondii* and other apicomplexans, rendering this protein redundant. WPDP contains a WIP-related protein domain which is known to facilitate actin binding. Numerous Golgi-resident actin-binding proteins have previously been identified in mammalian cells and the actin cytoskeleton has been shown to be critical for maintaining Golgi structure and facilitating vesicular trafficking (75). Similar to Sec16-L, WPDP had a modest phenotype upon conditional knockdown. It is possible that WPDP may be one of several actin-binding proteins that help maintain Golgi structure in *T. gondii* and related apicomplexans.

We additionally identified four novel parasite-specific Golgi proteins, GLP1-4, which were all found to be non-essential. GLP3, which we identified as an interactor of ULP1 by IP, was found to localize upstream of GRASP55 like Sec16-L and Sec31, and therefore could play a role in ER to Golgi vesicle transport. GLP1 and GLP4 both localize downstream of GRASP55, suggesting they function in the trans-Golgi network. Both proteins exhibit only a relatively small reduction in plaque size upon their depletion. Interestingly, the two proteins are similar in size and both contain a region with one larger coiled-coil domain closely followed by two shorter coiled-coil domains. While their sequence similarity does not suggest they are paralogs, their similar localizations, phenotypes, and structural compositions could indicate possible functional redundancy. Finally, GLP2 was found to localize to the cis-medial Golgi and caused a dramatic 91.8% reduction in plaque size when depleted. Despite this, we did not observe any obvious growth defects or morphological changes by IFA, suggesting that loss of GLP2 may affect either invasion or egress rather than replication. Interestingly, all of the GLPs contain several predicted coiled-coil domains and are only found in the cyst-forming coccidia. Coiled-coil domain-containing proteins such as the golgin proteins GM130, giantin, and CASP are known to act as tethering factors in other eukaryotes (76). As the GLPs are unique to the cyst-forming coccidia, it is possible they could use their CC domains to act as Golgi tethering factors that carry out unique functions in this subset of apicomplexan parasites.

Together, this study reveals a total of 12 novel Golgi-associated proteins including several that are essential and others that are parasite-specific. Their identification provides valuable new markers for the secretory pathway and provides exciting candidates for further functional analysis. Due to the apicomplexan-specific nature of several of these proteins, they represent both novel cell biology and potential targets for therapeutic intervention.

## Materials and Methods

### *T. gondii* and host cell culture

Parental *T. gondii* RHΔ*hxgprtΔku80* (wild-type) and subsequent strains were grown on confluent monolayers of human foreskin fibroblasts (BJ, ATCC, Manassas, VA) at 37°C and 5% CO2 in Dulbecco’s Modified Eagle Medium (DMEM) supplemented with 5% fetal bovine serum (Gibco), 5% Cosmic calf serum (Hyclone), and 1x penicillin-streptomycin-L-glutamine (Gibco). Constructs containing selectable markers were selected using 1 μM pyrimethamine (dihydrofolate reductase-thymidylate synthase [DHFR-TS]), 50 μg/mL mycophenolic acid-xanthine (HXGPRT), or 40 μM chloramphenicol (CAT) (77–79). For AID conditional knockdown experiments, media was supplemented with 500 μM indoleacetic acid (Sigma-Aldrich; I2886) or vehicle control.

### Antibodies

The HA epitope was detected with mouse monoclonal antibody (mAb) HA.11 (BioLegend; 901515). The V5 epitope was detected with mouse mAb anti-V5 (Invitrogen; R96025). *Toxoplasma*-specific antibodies include rabbit pAb anti-IMC6 (80), rat pAb anti-DrpB (23), guinea pig pAb anti-NHE3 (81), mouse pAb anti-SERCA (82), mouse mAb anti-ROP7 (1B10) (83), mouse mAb anti-MIC2 (6D10) (84), mouse pAb anti-GRA14 (83), mouse mAb anti-ATrx1 (11G8) (85), mouse mAb anti-F1β subunit (5F4) (86), and mouse pAb anti-ROP13 (87).

### Endogenous epitope tagging and knockout

For C-terminal endogenous tagging, a pU6-Universal plasmid containing a protospacer against the 3′ untranslated region (UTR) of the target protein approximately 100 bp downstream of the stop codon was generated, as described previously (34). A homology-directed repair (HDR) template was PCR amplified using the Δ*ku80*-dependent LIC vector pmAID3xHA.LIC-HPT, pmIAA73xHA.LIC-HPT, p3xHA.LIC-DHFR, p3xV5.LIC-DHFR, or pTurboID3xHA.LIC-DHFR, all of which include the epitope tag, 3′ UTR, and a selection cassette (88). The 60-bp primers include 40 bp of homology immediately upstream of the stop codon or 40 bp of homology within the 3′ UTR downstream of the CRISPR/Cas9 cut site. For knockout of ULP1, the protospacer was designed to target the coding region of ULP1, ligated into the pU6-Universal plasmid and prepared similarly to the endogenous tagging constructs. The HDR template was PCR amplified from a pJET vector containing the HXGPRT drug marker driven by the NcGRA7 promoter using primers that included 40 bp of homology immediately upstream of the start codon or 40 bp of homology downstream of the region used for homologous recombination for endogenous tagging. All primers that were used for pU6-Universal plasmids and HDR templates are listed in Table S3.

For all tagging and knockout constructs, approximately 50 µg of the sequence-verified pU6-Universal plasmid was precipitated in ethanol, and the PCR-amplified HDR template was purified by phenol chloroform extraction and precipitated in ethanol. Both constructs were electroporated into the appropriate parasite strain. Transfected cells were allowed to invade a confluent monolayer of HFFs overnight, and appropriate selection was subsequently applied. Successful tagging was confirmed by IFA, and clonal lines of tagged parasites were obtained through limiting dilution.

### Immunofluorescence assay

Confluent HFF cells were grown on glass coverslips and infected with *T*. *gondii*. After 24 hours, the coverslips were fixed with 3.7% formaldehyde in PBS and processed for immunofluorescence as described (87). Primary antibodies were detected by species-specific secondary antibodies conjugated to Alexa Fluor 594/488 (ThermoFisher). Coverslips were mounted in Vectashield (Vector Labs), viewed with an Axio Imager.Z1 fluorescent microscope and processed with ZEN 2.3 software (Zeiss).

### Western blot

Parasites were lysed in 1x Laemmli sample buffer with 100 mM DTT and boiled at 100°C for 5 minutes. Lysates were resolved by SDS-PAGE and transferred to nitrocellulose membranes, and proteins were detected with the appropriate primary antibody and corresponding secondary antibody conjugated to horseradish peroxidase. Chemiluminescence was induced using the SuperSignal West Pico substrate (Pierce) and imaged on a ChemiDoc XRS+ (Bio-Rad).

### Plaque assay

HFF monolayers were infected with 200 parasites per well of individual strains (-/+ IAA for AID strains) and allowed to form plaques for 7 days. Cells were then fixed with ice-cold methanol and stained with crystal violet. To quantify plaque size, the areas of 50 plaques per condition were measured using ZEN software (Zeiss). To quantify plaque number, the total number of plaques in each condition was counted manually. All plaque assays were performed in triplicate. Graphical and statistical analyses were performed using Prism GraphPad 8.0.

### Quantification of parasites per vacuole

Parasites were pre-treated with IAA or vehicle control for 18 hours prior to infecting coverslips. After coverslips were infected, parasites were allowed to grow -/+ IAA for 30 hours prior to fixation and staining with mouse anti-HA (detecting ULP1^AID^) and rabbit anti-IMC6. For each condition, the number of parasites per vacuole (1, 2, 4, 8, 16, or ≥32) was counted for >100 vacuoles across 10 different fields. Experiments were performed in triplicate. Significance was determined using multiple two-tailed t tests.

### Affinity capture of biotinylated proteins

For affinity capture of proteins from whole cell lysates, HFF monolayers infected with ULP1^TurboID^ or control parasites (RHΔ*hxgprt*Δ*ku80*, WT) were grown in medium containing 150 µM biotin for 30 hours. Intracellular parasites in large vacuoles were collected by manual scraping, washed in PBS, and lysed in radioimmunoprecipitation assay (RIPA) buffer (50 mM Tris [pH 7.5], 150 mM NaCl, 0.1% SDS, 0.5% sodium deoxycholate, 1% NP-40) supplemented with Complete Protease Inhibitor Cocktail (Roche) for 30 min on ice. Lysates were centrifuged for 15 min at 14,000 x g to pellet insoluble material, and the supernatant was incubated with Streptavidin Plus UltraLink resin (Pierce) overnight at 4°C under gentle agitation. Beads were collected and washed five times in RIPA buffer, followed by three washes in 8 M urea buffer (50 mM Tris-HCl [pH 7.4], 150 mM NaCl) (89). Samples were submitted for on-bead digests and subsequently analyzed by mass spectrometry. The experiment was performed in duplicate.

### Immunoprecipitation assay

ULP1^3XHA^ was isolated from 5×10^9^ *T. gondii* RH tachyzoites lysed in NP-40 lysis buffer (1% NP-40, 150 mM NaCl, 50 mM Tris [pH 8.0]) supplemented with complete protease inhibitor (Roche). The insoluble material was removed from the lysate by centrifugation at 10,000 x *g* for 20 minutes. The soluble lysate fraction was incubated with rat anti-HA affinity matrix (Roche) for 3 hours at room temperature with gentle agitation. After washing in NP-40 lysis buffer, the bound protein was eluted using high pH (100 mM triethylamine, pH 11.5), dried to a pellet, and submitted for mass spectrometry.

### Mass spectrometry

Samples were resuspended in digestion buffer (8M Urea, 0.1M Tris-HCl [pH 8.5]) and then were reduced, alkylated, and digested by sequential addition of Lys-C and trypsin proteases. Samples were then desalted using C18 tips (Pierce) and fractionated online using a 75-μm inner-diameter fritted fused silica capillary column with a 5-μm pulled electrospray tip and packed in-house with 25 cm of C18 (Dr. Maisch GmbH) 1.9-μm reversed-phase particles. The gradient was delivered by a 140-minute gradient of increasing acetonitrile and eluted directly into a Thermo Orbitrap Fusion Lumos instrument where MS/MS spectra were acquired by Data Dependent Acquisition (DDA). Data analysis was performed using ProLuCID and DTASelect2 implemented in Integrated Proteomics Pipeline IP2 (Integrated Proteomics Applications) (90–92). Database searching was performed using a FASTA protein database containing *T*. *gondii* GT1-translated open reading frames downloaded from ToxoDB. Protein and peptide identifications were filtered using DTASelect and required a minimum of two unique peptides per protein and a peptide-level false positive rate of less than 5% as estimated by a decoy database strategy. Candidates were ranked by spectral count comparing experimental versus control samples (93). TurboID results were filtered to include only proteins that were at least two-fold enriched with a difference of >5 spectral counts when comparing ULP1^TurboID^ samples versus control samples. IP results were filtered to include only proteins that were at least two-fold enriched when comparing ULP1^3xHA^ IP samples versus control samples.

### Bioinformatic analysis of novel proteins

Functional domains were identified using a combination of Phyre2, Interpro Domains, PFAM, and Panther analysis (28,30,94,95). Coiled coil domains were predicted using DeepCoil2 using a probability cut-off of 0.5 (31). Orthology was determined using OrthoMCL and BLASTp search against the VEuPathDB database (46).

## Acknowledgements

We thank Vern Carruthers for supplying the anti-MIC2 antibody, Gustavo Arrizabalaga for supplying the anti-NHE3 antibody, and David Sibley for supplying the anti-SERCA antibody. This work was supported by NIH grants AI123360 to P.J.B, and GM089778 to J.A.W. R.R.P. and V.Y. were supported by the Ruth L. Kirschstein National Research Service Award AI007323. R.R.P was additionally supported by the Ruth L. Kirschstein National Research Service Award GM007185 and the UCLA Molecular Biology Institute Whitcome Fellowship. J.J.Q was supported by a Milton Gottlieb endowment award and by a Beckman Scholars Award through the Arnold and Mabel Beckman Foundation.

---

**Table S1. ULP1 TurboID results.**

List of genes identified by mass spectrometry in the ULP1 TurboID experiment. Spectral counts are shown for each gene. “Enrichment Diff” refers to the difference between the average spectral count in ULP1^TurboID^ and control parasites. “Enrichment Fold” refers to the average spectral count for ULP1^TurboID^ samples divided by the average spectral count for control samples. GWCS = phenotype score assigned in a genome wide CRISPR/Cas9 screen (33). SP = signal peptide. #TMDs = number of transmembrane domains.

**Table S2. ULP1 immunoprecipitation results.**

List of genes identified by mass spectrometry in the ULP1 immunoprecipitation experiment. Spectral counts are shown for each gene. “Enrichment Diff” refers to the difference between the spectral count in ULP1^3xHA^ IP versus control experiments. “Enrichment Fold” refers to the spectral count for the ULP1^3xHA^ IP sample divided by the spectral count for the control sample. GWCS = phenotype score assigned in a genome wide CRISPR/Cas9 screen (33). SP = signal peptide. #TMDs = number of transmembrane domains.

**Table S3. Oligonucleotides used in this study.**

